# VAPA mediates lipid exchange between *Leishmania amazonensis* and host macrophages

**DOI:** 10.1101/2024.10.03.616432

**Authors:** Ilona Gdovinova, Albert Descoteaux

## Abstract

*Leishmania* is a vacuolar pathogen that replicates within parasitophorous vacuoles inside host phagocytes. To promote its replication, *Leishmania* relies on a panoply of strategies to acquire macromolecules such as lipids from host macrophages. In this study, we have evaluated the role of VAPA, an endoplasmic reticulum-resident membrane protein involved in inter-organellar lipid transport, in macrophages infected with *L. amazonensis*. Following infection of bone marrow-derived macrophages with metacyclic *L. amazonensis* promastigotes, we observed that VAPA gradually associates with communal parasitophorous vacuoles. Knockdown of VAPA prevented the replication of *L. amazonensis*, which was accompanied by an impaired parasitophorous vacuole expansion. Using fluorescent ceramide, we established that VAPA is required for the transport of sphingolipids to the parasitophorous vacuoles and for its acquisition by *L. amazonensis* amastigotes. Proximity-ligation and immunoprecipitation assays revealed that *L. amazonensis* hijacks VAPA by disrupting its interactions with the lipid transfer proteins CERT and ORP1L. Finally, we found that VAPA is essential for the transfer of the *Leishmania* virulence glycolipid lipophosphoglycan from the parasitophorous vacuoles to the host cell endoplasmic reticulum. We propose that VAPA contributes to the ability of *L. amazonensis* to colonize macrophages by mediating bi-directional transfer of lipids essential for parasite replication and virulence between the parasitophorous vacuoles and the host cell endoplasmic reticulum.

**AUTHOR SUMMARY:** The protozoan parasite *Leishmania amazonensis* replicates in macrophages, within communal parasitophorous vacuoles. To satisfy its various auxotrophies, this parasite must obtain macronutrients and metabolites from its host cell, including lipids. To salvage host sphingolipids, we obtained evidence that *L. amazonensis* exploits a macrophage nonvesicular lipid transport mechanism that requires the endoplasmic reticulum membrane protein VAPA. Moreover, we found that VAPA is also required for the transfer of the *Leishmania* virulence glycolipid lipophosphoglycan from the parasitophorous vacuole to the macrophage endoplasmic reticulum. The fact that VAPA is essential for *L. amazonensis* to colonize macrophages is consistent with the central role that VAPA plays in mediating bi-directional transfer of lipids and illustrates the importance of the host cell endoplasmic reticulum in this host-parasite interaction.

## INTRODUCTION

The protozoan parasite *Leishmania* is a vacuolar pathogen responsible for the leishmaniases, a spectrum of human diseases ranging from a confined cutaneous lesion to a progressive visceral infection that can be fatal if left untreated (1). Following inoculation into a mammalian host by an infected sand fly, infectious promastigote forms of the parasite are taken up by phagocytes. Internalized promastigotes evade microbicidal mechanisms associated to phagocytosis by subverting phagolysosomal biogenesis to generate parasitophorous vacuoles (PVs) in which they differentiate and replicate as mammalian stage amastigotes (2–9). To obtain the membrane required for expansion and maintenance of those PVs, *Leishmania* modulates their interactions with trafficking vesicles and sub-cellular compartments by hijacking the host cell membrane fusion machinery (10–16). Hence, knockdown or deletion of host cell soluble *N*-ethylmaleimide-sensitive factor attachment protein receptors (SNAREs) associated to the endoplasmic reticulum (ER) and to endosomes inhibited parasite replication and PV enlargement (10, 14, 17).

*Leishmania* displays various auxotrophies and must therefore acquire essential host-derived nutrients and macromolecules to promote its replication (18–20). In this context, lipid metabolism in *Leishmania* is of interest as it differs from that in mammalian cells and may be exploited for the development of novel therapeutic approaches (21, 22). Hence, in contrast to insect stage promastigotes which rely on *de novo* synthesis to produce the majority of their lipids, amastigotes acquire most of their lipids from the mammalian host (23). In the case of sphingolipids, evidence that *de novo* synthesis is unnecessary for the proliferation of intramacrophage amastigotes arose from the observation that *L. major* mutants defective in ceramide and sphingolipid biosynthesis were fully infective in an experimental model of cutaneous leishmaniasis (24, 25). Rather, *Leishmania* amastigotes acquire and remodel host sphingolipids, which contributes to their ability to colonize macrophages (26). However, the mechanism by which amastigotes salvage mammalian host sphingolipids remains to be elucidated.

In mammalian cells the ER is the site of *de novo* synthesis of ceramide, the precursor of most sphingolipids (27). Ceramide is transported to the Golgi where it is converted to sphingomyelin and other complex sphingolipids (28). Transfer of ceramide from the ER to the Golgi is mediated by the ceramide transport protein CERT which associates with the ER-resident membrane protein VAMP-associated proteins (VAPs) (29). Those proteins are components of membrane contact sites (MCS), which are regions of close contact between organelles specialized in the non-vesicular trafficking of molecules such as lipids and ions between organelles (30–32). In recent years, MCS components were shown to be hijacked by vacuolar pathogens to establish MCS between pathogen-containing vacuoles and host cell organelles, as a strategy to promote their intracellular replication (33–36). One example is the bacterial pathogen *Chlamydia trachomatis*, which through the action of secreted effectors recruits CERT and other host factors to the inclusion and brings the ER in close proximity (37–39). Through interactions with VAPA/B, *C. trachomatis* then reroutes sphingolipid biosynthesis to its replicative niche (38, 40). Given the importance of the ER in the synthesis and inter-organelle trafficking of lipids, we explored the role of VAPA in the context of *L. amazonensis*-infected macrophages. Our findings indicate that *L. amazonensis* amastigotes exploits VAPA to promote their replication through the bi-directional exchange of lipids essential for parasite replication and virulence. This study highlights the importance of the host cell ER for the intracellular replication of *L. amazonensis*.

## RESULTS

### The ER protein VAPA associates with *L. amazonensis*-harboring communal parasitophorous vacuoles

Previous studies revealed that *L. amazonensis*-harboring communal PVs are hybrid compartments composed of both host ER and endocytic pathway components (10, 41). Those PVs acquire ER content and membrane components through continuous interactions with this organelle (41). The protein VAPA is a prominent ER membrane theter involved in the interactions and lipid transfer between the ER and other organelles through the formation of MCS (42). To investigate the potential role of VAPA in the intracellular fate of *L. amazonensis*, we first determined the kinetics of its association with PVs. To this end, we infected bone marrow-derived macrophages (BMM) with *L. amazonensis* metacyclic promastigotes and we used confocal immunofluorescence microscopy to assess the fate of VAPA at various time points post-phagocytosis. As controls, we infected BMM with *L. major* metacyclic promastigotes, which differentiate and replicate as amastigotes in tight-fitting individual PVs and we also fed BMM with serum-opsonized zymosan. As illustrated in Figure 1A-B we found that VAPA accumulates on communal PVs containing *L. amazonensis* amastigotes starting at 24 h post-phagocytosis. In contrast, we observed that VAPA was associated with a significantly smaller subset of PVs harboring *L. major* amastigotes and of phagosomes containing zymosan. Western blot analysis revealed no variations in VAPA levels in *L. amazonensis*-infected BMM over a 72 h post-infection kinetics (Fig 1D). To provide further evidence that VAPA associates with the membrane of communal PVs, we assessed its colocalization with LAMP1 which is also recruited to these PVs. As shown in Figure 1C, VAPA colocalizes with LAMP1 at the PV membrane. These results indicate that VAPA associates with communal PVs containing *L. amazonensis* amastigotes and raise the possibility that it contributes to their development.

**Figure 1.**
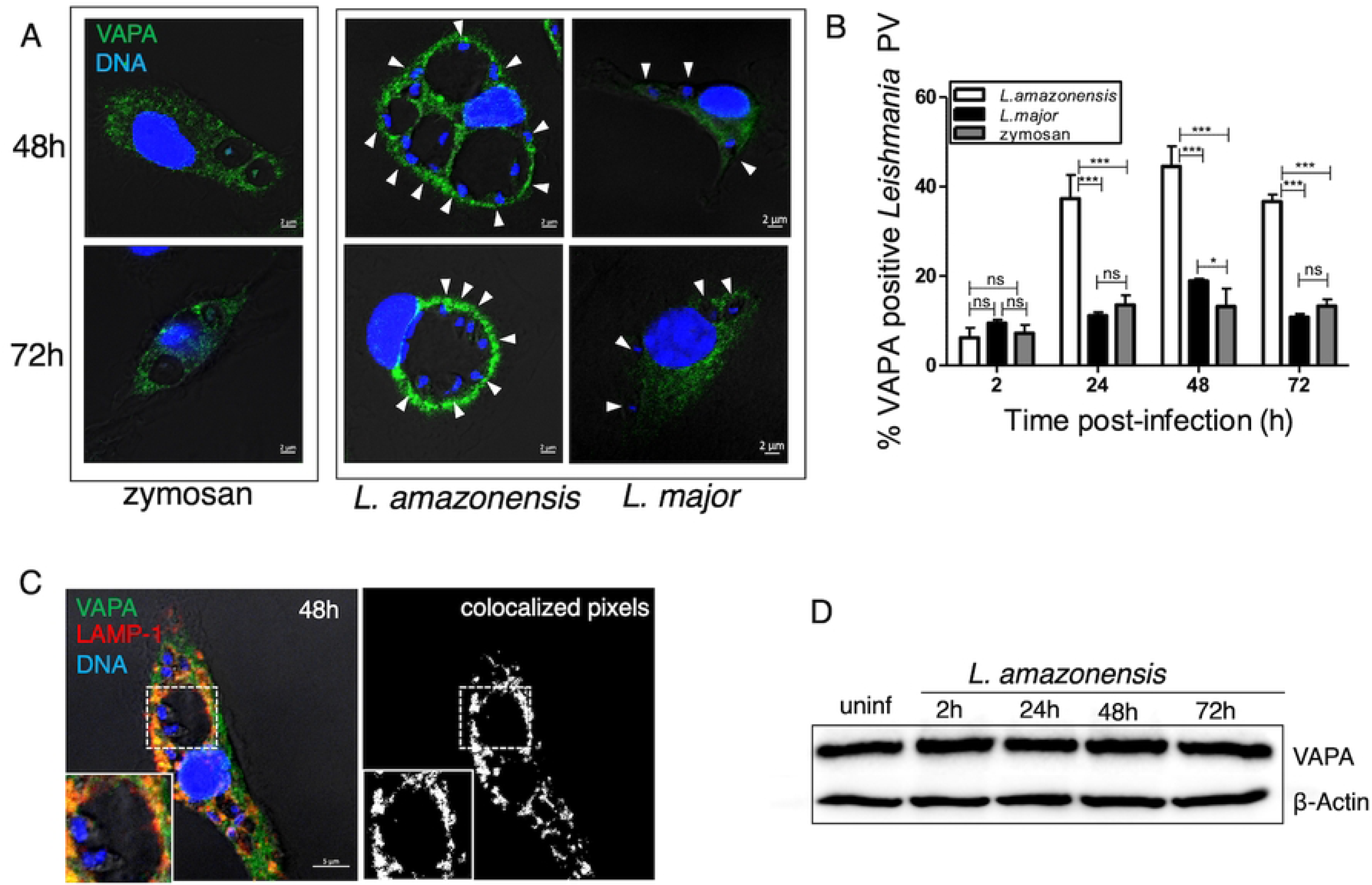
VAPA associates with *L. amazonensis*-harboring communal PVs. BMM were infected with *L .amazonensis* or *L. major* metacyclic promastigotes for the indicated time points. Controls consisted of BMM fed with zymosan. (**A**) Distribution of VAPA (green) and its association with PVs were assessed by confocal immunofluorescence microscopy. DNA is shown in blue. (**B**) Quantification of the association of VAPA to *L. amazonensis-* and *L. major*-harboring PVs and to zymosan-containing phagosomes. (**C**) Colocalization of LAMP1 with VAPA on communal PVs harboring *L.amazonensis* at 48 h post-infection. Colocalization (white pixels) of LAMP-1 (red) with VAPA (green) was assessed and quantified by confocal immunofluorescence microscopy. Insets display the PV area. DNA is shown in blue. (**D**) Western blot analysis of VAPA levels over time in *L. amazonensis*-infected BMM. Data are presented as the means ± standard errors of the means (SEM) of values from three independent experiments. White arrowheads denote internalized parasites. Representative images from three experiments are shown. **, *P* ≤ 0.01; ***, *P* ≤ 0.001.

### VAPA contributes to the replication of *L. amazonensis* and PV expansion

Given the association of VAPA with communal PVs, we sought to investigate its role in the development of those compartments and on the intracellular fate of *L. amazonensis*. We infected BMM (untreated or treated with either siRNA to VAPA or scrambled siRNA) with *L. amazonensis* metacyclic promastigotes and at various time points post-infection we assessed parasite burden and PV surface area. We also included BMM infected with *L. major* metacyclic promastigotes in our analyses. As depicted in Figure 2A, whereas VAPA knockdown had no impact on the phagocytosis of *L. amazonensis*, it severely restricted parasite replication. The inability of *L. amazonensis* to replicate in VAPA knockdown BMM correlated with smaller PVs size at 48 h and 72 h post-infection (Fig 2B). In the case of *L. major*, we noted a slight reduction in its replication in VAPA knockdown BMM at 24 h and 48 h post-infection and at 72 h post-infection we did not detect a significant impact of VAPA knockdown on parasite replication (Fig 2C). These results indicate a role for VAPA in the replication of *L. amazonensis* amastigotes and in communal PV expansion.

**Figure 2.**
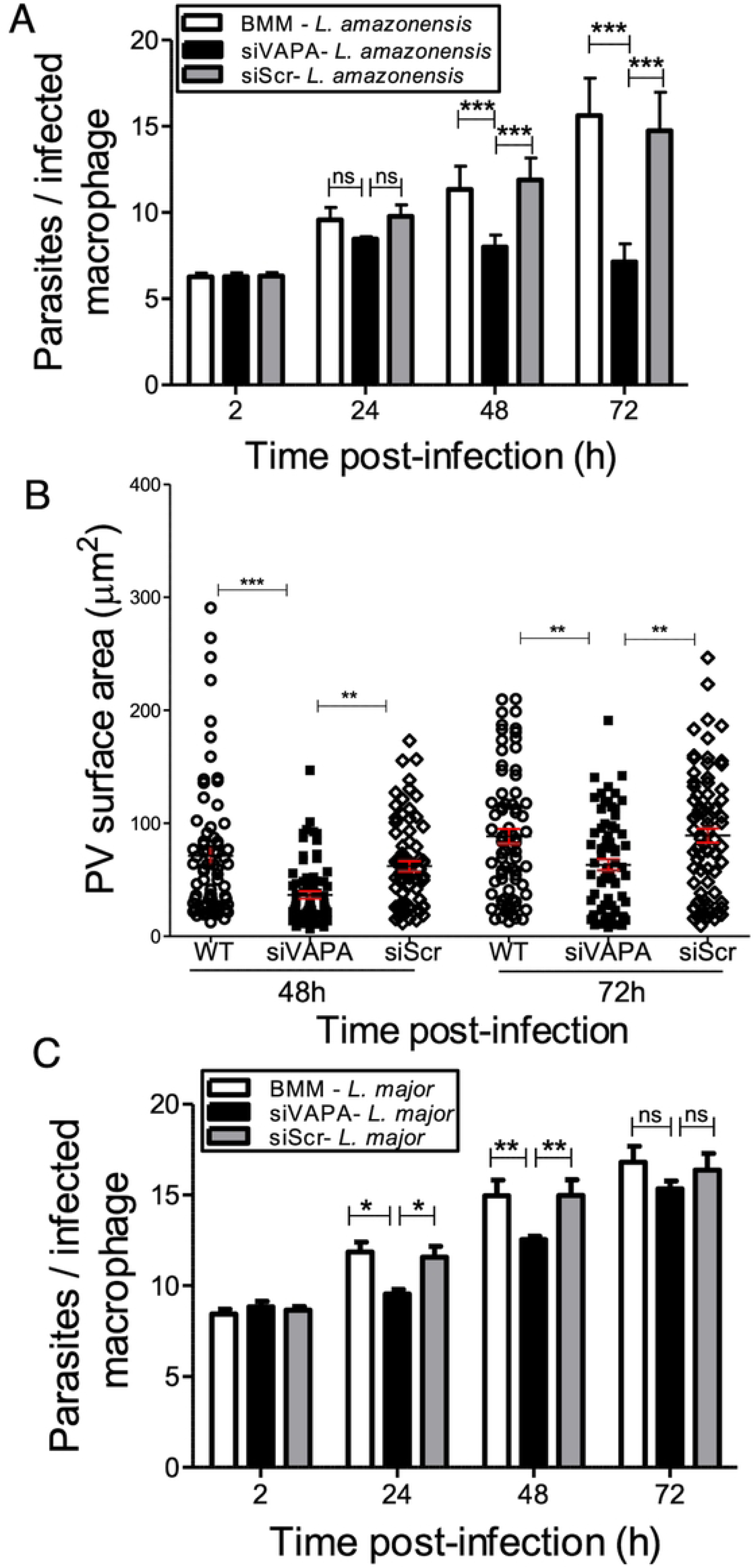
VAPA regulates the replication of *L. amazonensis* and PV expansion. BMM (untreated, treated with siRNA, or treated with scrambled siRNA) were infected with *L. amazonensis* or *L. major* metacyclic promastigotes and at various time points post-phagocytosis parasite replication and PV size were assessed. (**A**) Quantification of *L. amazonensis* parasite burden at 2, 24, 48, and 72 h post-infection. Data are presented as the means ± SEM of values from three independent experiments. **, *P* ≤ 0.01; ***, *P* ≤ 0.001. (**B**) Quantification of PV size in BMM infected with *L. amazonensis* at 48 and 72 h post-phagocytosis. Data are presented as a cloud with means ± standard deviations (SD) of values from three independent experiments for a total of 450 PVs. **, *P* ≤ 0.01; ***, *P* ≤ 0.001. (**C**) Quantification of *L. major* parasite burden at 2, 24, 48, and 72 h post-infection. Data are presented as the means ± SEM of values from three independent experiments. *, *P* ≤ 0.05; **, *P* ≤ 0.01.

### Acquisition of sphingolipids by *L. amazonensis* requires VAPA

*Leishmania* amastigotes salvage most of their lipids, including sphingolipids, from the host (23, 25). Since VAPA is involved in the transfer of lipids synthesized in the ER to their destination organelle (43), we assessed whether it mediates the transfer of ceramide/sphingolipids to the PVs. To this end, BMM (untreated, treated with either siRNA to VAPA or scrambled siRNA, or mock lipofected) infected for 48 h with *L. amazonensis* metacyclic promastigotes were labeled with fluorescent ceramide (BODIPY FL C5-Ceramide). Using live cell imaging, we observed incorporation of fluorescent sphingolipids into *L. amazonensis* amastigotes present within PVs in control groups (Figure 3A-B). In contrast, *L. amazonensis* amastigotes did not acquire fluorescent sphingolipids in VAPA knockdown BMM, consistent with a role for VAPA in the acquisition of host sphingolipids by *L. amazonensis*. VAPA was previously shown to be involved in the transfer of ceramide/sphingolipids to the *Chlamydia* inclusion through interactions with the ceramide transfer protein CERT (37, 38). We therefore sought to determine whether VAPA associates with CERT and contributes to the acquisition of ceramide/sphingolipids by *L. amazonensis*. We included as a control the endosomal/lysosomal oxysterol lipid binding protein ORP1L which also associates with VAPA (44). Following phagocytosis of *L. amazonensis* metacyclic promastigotes by BMM, we observed by immunofluorescence confocal microscopy that the distribution of CERT and ORP1L was uniform at 2 h and 24 h post-phagocytosis. At 72 h post-phagocytosis, we found that labeling for CERT and to a lesser extent ORP1L, was more intense around communal PVs and both were partly co-localizing with VAPA (Figure 4 A-B). Given that both CERT and ORP1L associate with *L. amazonensis*-harboring PVs, we next assessed the potential role of CERT and ORP1L on the replication of *L. amazonensis* and on PV size. As illustrated in Fig 4 C-D, CERT knockdown had a minimal impact on the replication of *L. amazonensis* but resulted in a reduced PV size. In contrast, ORP1L knockdown resulted in increased parasitemia despite a defect in PV expansion (Fig 4 E-F).

**Figure 3.**
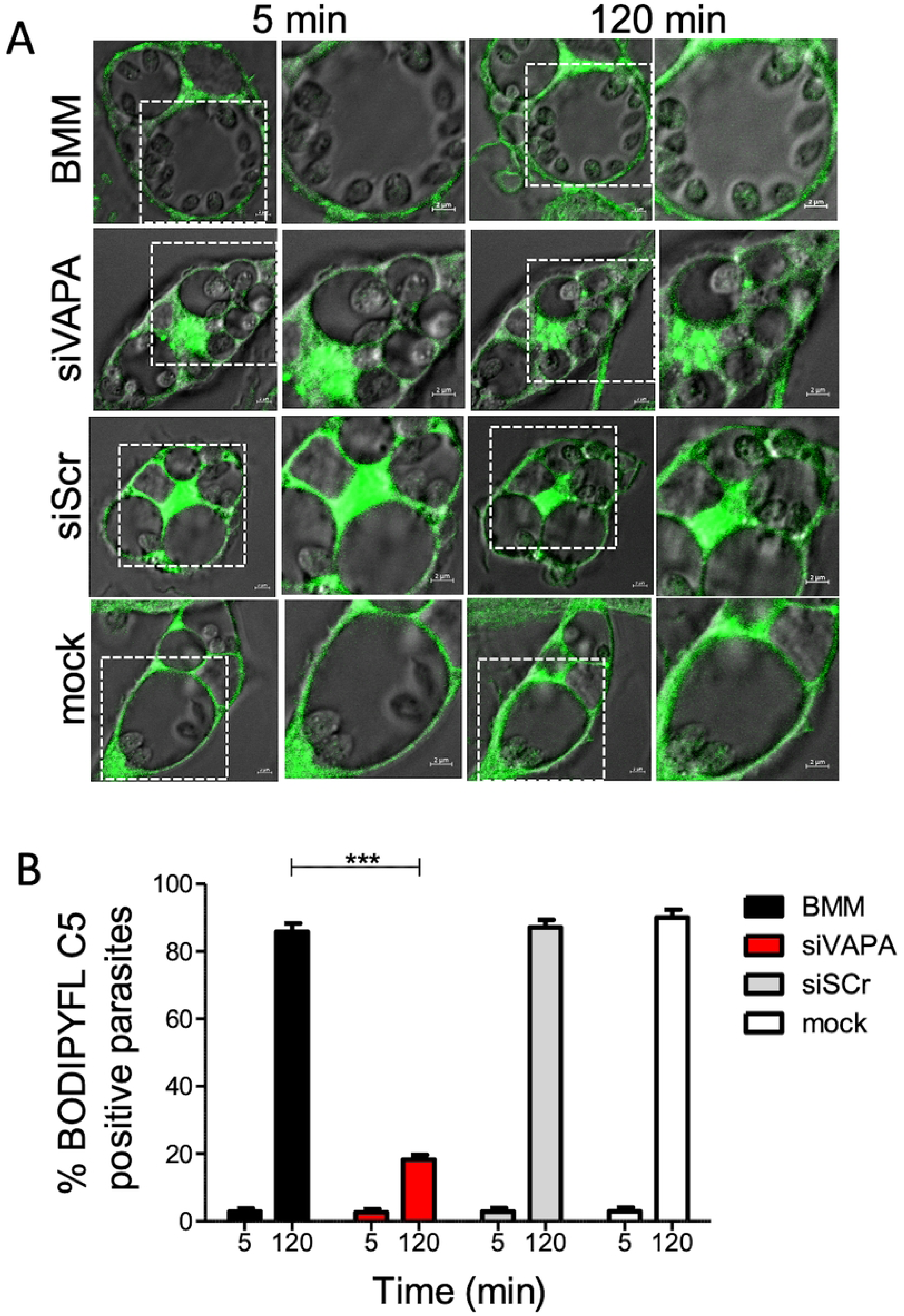
VAPA is required for the acquisition of sphingolipids by *L. amazonensis*. BMM (untreated, treated with siRNA to VAPA, treated with scrambled siRNA, and mock transfected) were infected with *L. amazonensis* metacyclic promastigotes and at 48 h post-infection BMM were incubated with BODIPY FL-C5-Ceramide (green). (**A**) Images of live cells were acquired at 5 min and 120 min after the addition of BODIPY FL C5-Ceramide. (**B**) Quantification of BODIPYFL-C5 positive *L. amazonensis* amastigotes at 5 min and 120 min after the addition of BODIPY FL-C5-Ceramide. Data are presented with means ± standard deviations (SD) of values from three independent experiments. ***, *P* ≤ 0.001 (in comparison to controls, siScr and mock).

**Figure 4.**
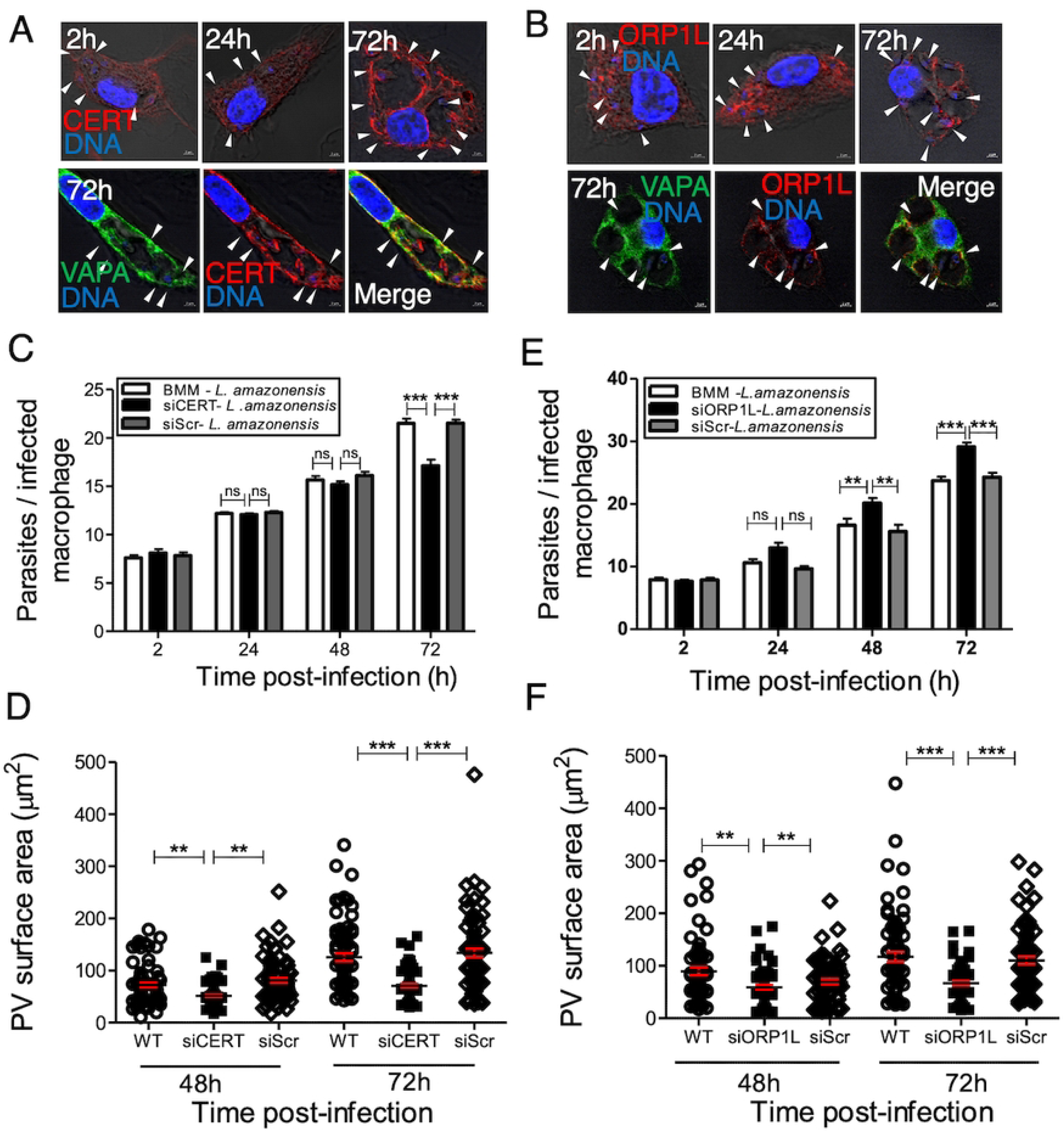
CERT and ORP1L associate with *L. amazonensis*-harboring communal PVs and regulate their expansion. BMM were infected with *L. amazonensis* metacyclic promastigotes and at the indicated time points the localization of CERT (red) (**A**) and ORP1L (red) (**B**) was assessed by immunofluorescence confocal microscopy. Co-localization of VAPA (green) with CERT (red) (**A**) and ORP1L(red) (**B**) was assessed by immunofluorescence confocal microscopy at 72 h post-infection. DNA is shown in blue. White arrowheads denote internalized parasites. Representative images from three independent experiments are shown. BMM (normal, treated with siRNA to CERT or scramble siRNA) were infected with *L. amazonensis* metacyclic promastigotes and at the indicated time points post-infection, parasite burden (**C**) and PV size (**D**) were assessed. BMM (untreated, treated with siRNA to ORP1L or scramble siRNA) were infected with *L. amazonensis* metacyclic promastigotes and at the indicated time points post-infection parasite burden (**E**) and PV size (**F**) were assessed. Data are presented as the means ± SD of values from three independent experiments. **, *P* ≤ 0.01; ***, *P* ≤ 0.001. For the determination of PV surface area (**D**, **F**) data are presented as clouds with means ± SD of values from three independent experiments for a total of 450 PVs. **, *P* ≤ 0.01; ***, *P* ≤ 0.001.

### *L. amazonensis* disrupts VAPA-CERT and VAPA-ORP1L

In the context of membrane contact sites, co-localization by immunofluorescence confocal microscopy may not be sufficient to conclude that VAPA forms complexes with CERT and ORP1L. We therefore used proximity ligation assay (PLA) to assess *in situ* VAPA-CERT and VAPA-ORP1L protein-protein interactions in uninfected BMM and in BMM infected with *L. amazonensis* metacyclic promastigotes for 24 h, 48 h, and 72 h. With PLA, proximity between two proteins (below 40 nm) is detected by fluorescent dots. As shown in Figure 5A-D, VAPA forms complexes with CERT and ORP1L in uninfected BMM. Remarkably, both VAPA-CERT and VAPA-ORP1L *in situ* complexes disappeared within 6 h post-infection in infected BMM, indicating that *L. amazonensis* efficiently disrupts these interactions. Previous studies revealed that insertion of the *Leishmania* pathogenicity glycolipid LPG into host cell membranes disrupts lipid microdomains, thereby altering protein distribution (7, 45, 46). Similarly, the *Leishmania* surface metalloprotease GP63 targets and cleaves a subset of host cell proteins, which may affect protein complex formation (47). To investigate the possibility that either LPG or GP63 are involved in the disruption of *in situ* VAPA-ORP1L complexes, we infected BMM with Δ*lpg1* and Δ*lpg1*+*LPG1 L. donovani* and with Δ*gp63* and Δ*gp63*+*GP63 L. major* metacyclic promastigotes. At 6 h and 24 h post-infection, we assessed VAPA-ORP1L *in situ* interactions through PLA. VAPA-ORP1L complexes were disrupted at 6 h post-infection regardless of the presence or absence of LPG and GP63 (Figure 5E-F). We noted that infection of BMM with *L. major* and *L. donovani*, which replicate in tight-fitting individual PVs, did not fully disrupt VAPA-ORP1L complexes as observed with *L. amazonensis*. At 24 h post-infection, we observed a significant increase in the number of VAPA-ORP1L *in situ* complexes in BMM infected with the Δ*gp63 L. major* mutant with respect to 6 h post-infection time point (Fig 5F). These results suggest that *L. amazonensis* hijacks VAPA and that neither LPG nor GP63 are responsible for the disruption of VAPA-CERT and VAPA-ORP1L. GP63 might however play a role in preventing the formation of those complexes at later time point post-infection.

**Figure 5.**
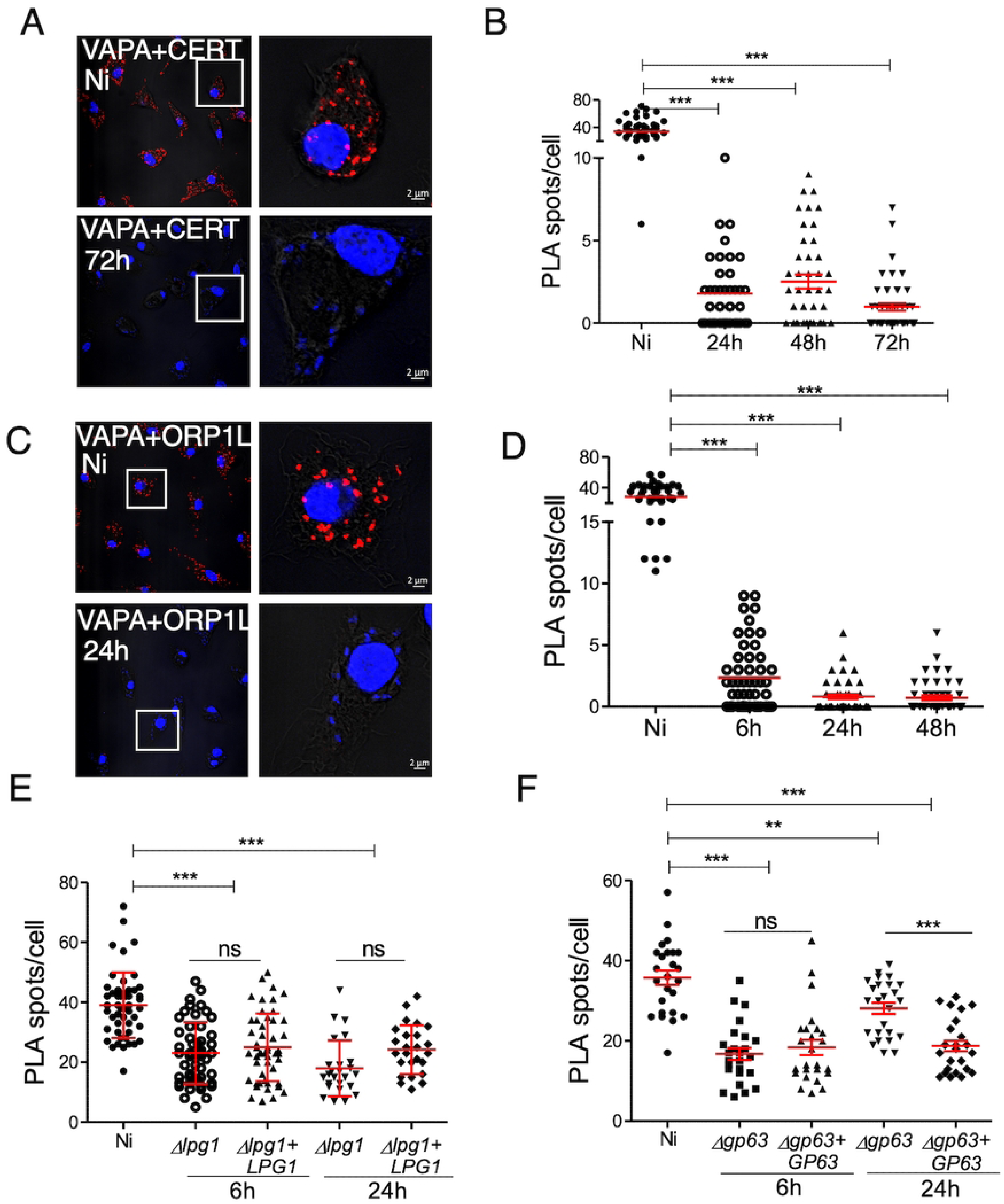
*L. amazonensis* hijacks VAPA. BMMs were infected or not with either *L. amazonensis* (**A-D**), Δ*lpg1* and Δ*lpg1* + *LPG1 L. donovani* (**E**) or *Δgp63* and *Δgp63*+*GP63 L*. *major* (**F**) metacyclic promastigotes. At the indicated time points post-infection, *in situ* VAPA-CERT (**A**) and VAPA-ORP1L (**C**) complexes with detected by proximity ligation and visualized by confocal immunofluorescence microscopy (red dots). DNA is in blue. Representative images from 3 independent experiments are shown. Quantification of *in situ* complexes for VAPA-CERT (**B**) and VAPA-ORP1L (**D**) in uninfected BMM and in BMM infected with *L. amazonensis*. Quantification of *in situ* complexes for VAPA-ORP1L in uninfected BMM and in BMM infected with either (**E**) Δ*lpg1* and Δ*lpg1* + *LPG1 L. donovani* or (**F**) *Δgp63* and *Δgp63*+*GP63 L*. *major*. Data are presented as clouds with means ± standard deviations (SD) of values from three independent experiments for a total of 75 cells in each group. **, *P* ≤ 0.01; ***, *P* ≤ 0.001.

### VAPA is required for the transfer of LPG out of the PV

LPG, the major lipid component of the *Leishmania* promastigote surface coat (48), is released from the parasite following phagocytosis and redistributes to the ER of infected macrophages (49). We previously reported that trafficking out of the PVs of this *Leishmania* virulence glycolipid requires the SNAREs Sec22b and syntaxin-5, which regulate ER-Golgi trafficking (49). Given that VAPA participates in multiple lipid transport pathways from the ER to destination organelles (43, 50), we investigated the possibility that this protein contributes to the transfer of LPG from the PV to the ER. First, we infected BMM with *L. major* metacyclic promastigotes and at 6 h post-phagocytosis, we analyzed the distribution of LPG with respect to VAPA by confocal immunofluorescence microscopy. We used *L. major* instead of *L. amazonensis* because the anti-LPG antibody CA7AE recognizes more efficiently *L. major* LPG. As shown in Figure 6A, LPG released by *L. major* promastigotes co-localized with VAPA, suggesting a possible interaction between these two molecules. Next, we infected BMM (untreated, treated with either siRNA to VAPA or scrambled siRNA, mock transfected) with *L. major* metacyclic promastigotes and at 6 h post-phagocytosis we assessed the distribution of LPG. The distribution of LPG within infected BMM was strikingly altered in the absence of VAPA, as depicted in Figure 6B. Hence, whereas LPG was evenly distributed throughout control infected BMM, it was confined to the cell periphery in VAPA knockdown BMM. Further analysis indicated that in the absence of VAPA, LPG was mainly associated with late endosomes/lysosomes, as it co-localizes with LAMP1 Figure 6C. These results indicate that VAPA is involved in the transport of LPG from the PVs to the host cell ER and suggest that it can mediate bi-directional lipid transport.

**Figure 6.**
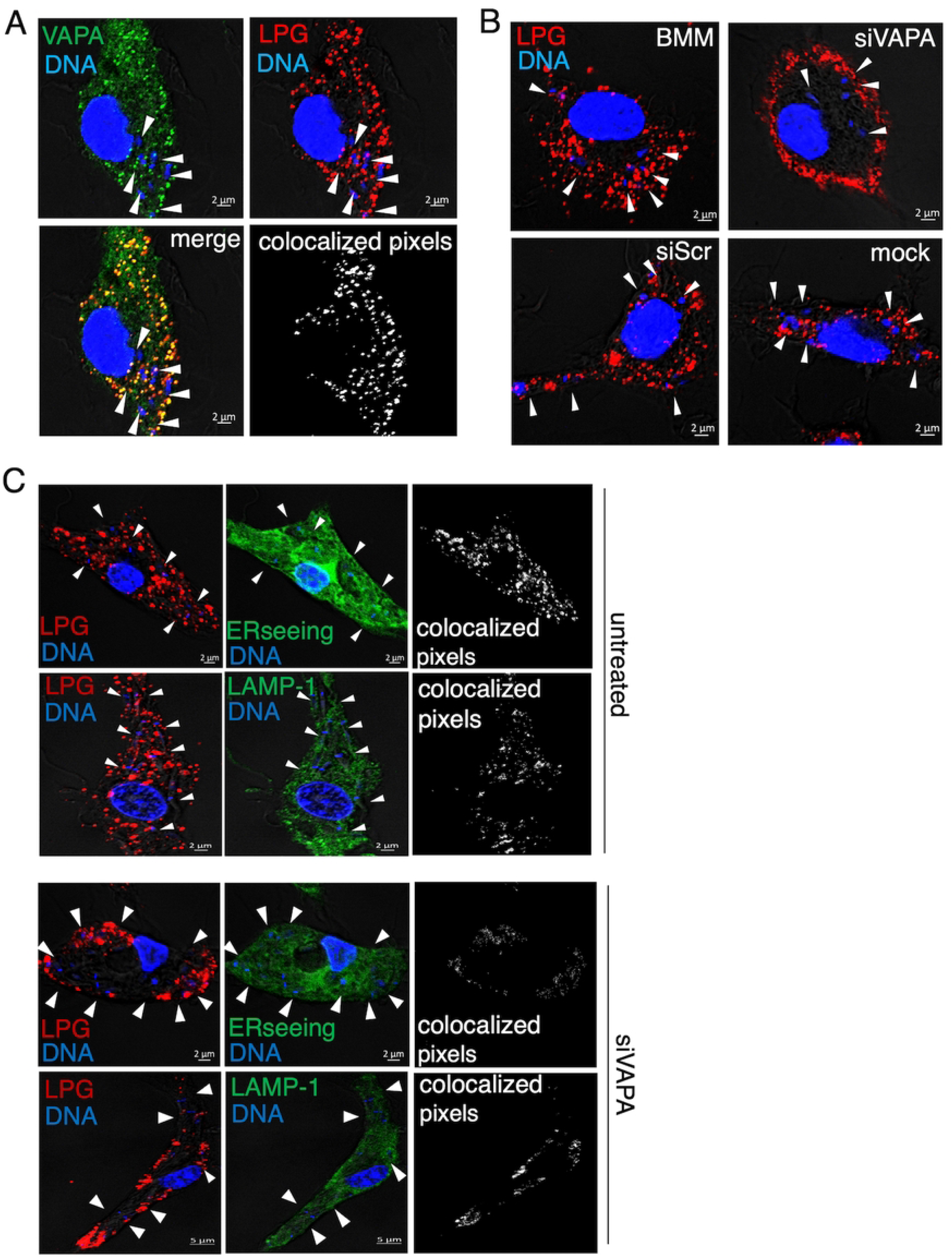
VAPA is required for the transfer of LPG to the host cell ER. (**A**) BMM were infected with *L. major* metacyclic promastigotes and at 6 h post-phagocytosis, the localization of VAPA (green) and LPG (red) were assessed by confocal immunofluorescence microscopy. Colocalized pixels are in white and DNA is in blue. (**B**) BMM (untreated or treated with siRNA to VAPA, scrambled siRNA, or mock transfected) were infected with *L. major* metacyclic promastigotes and at 6 h post-phagocytosis, the localization of LPG (red) was assessed by confocal immunofluorescence microscopy. DNA is in blue. (**C**). Untreated BMM (left panel) or BMM treated with siRNA to VAPA (right panel) were infected with *L. major* metacyclic promastigotes and at 6 h post-phagocytosis, the co-localization of LPG (red) with ERSeeing (green) or LAMP1 (green) was assessed by confocal immunofluorescence microscopy. Colocalized pixels are in white, DNA is in blue. White arrowheads denote internalized parasites. Representative images from 3 independent experiments are shown.

## DISCUSSION

Vacuolar pathogens scavenge macromolecules, nutrients and metabolites from their host cell. To this end, they establish communication between the vacuole in which they replicate and host cell organelles (33, 35, 36). In the present study, we investigated the role of the macrophage ER membrane protein VAPA in the interaction between *L. amazonensis* and its host cell. We found that VAPA is essential for *L. amazonensis* replication and PV expansion as well as for the acquisition of sphingolipids by the parasite. Moreover, we found that VAPA is essential for the transfer of the virulence glycolipid LPG from the PV to the host cell ER, indicating that VAPA mediates bi-directional lipid transport (Figure 7).

**Figure 7.**
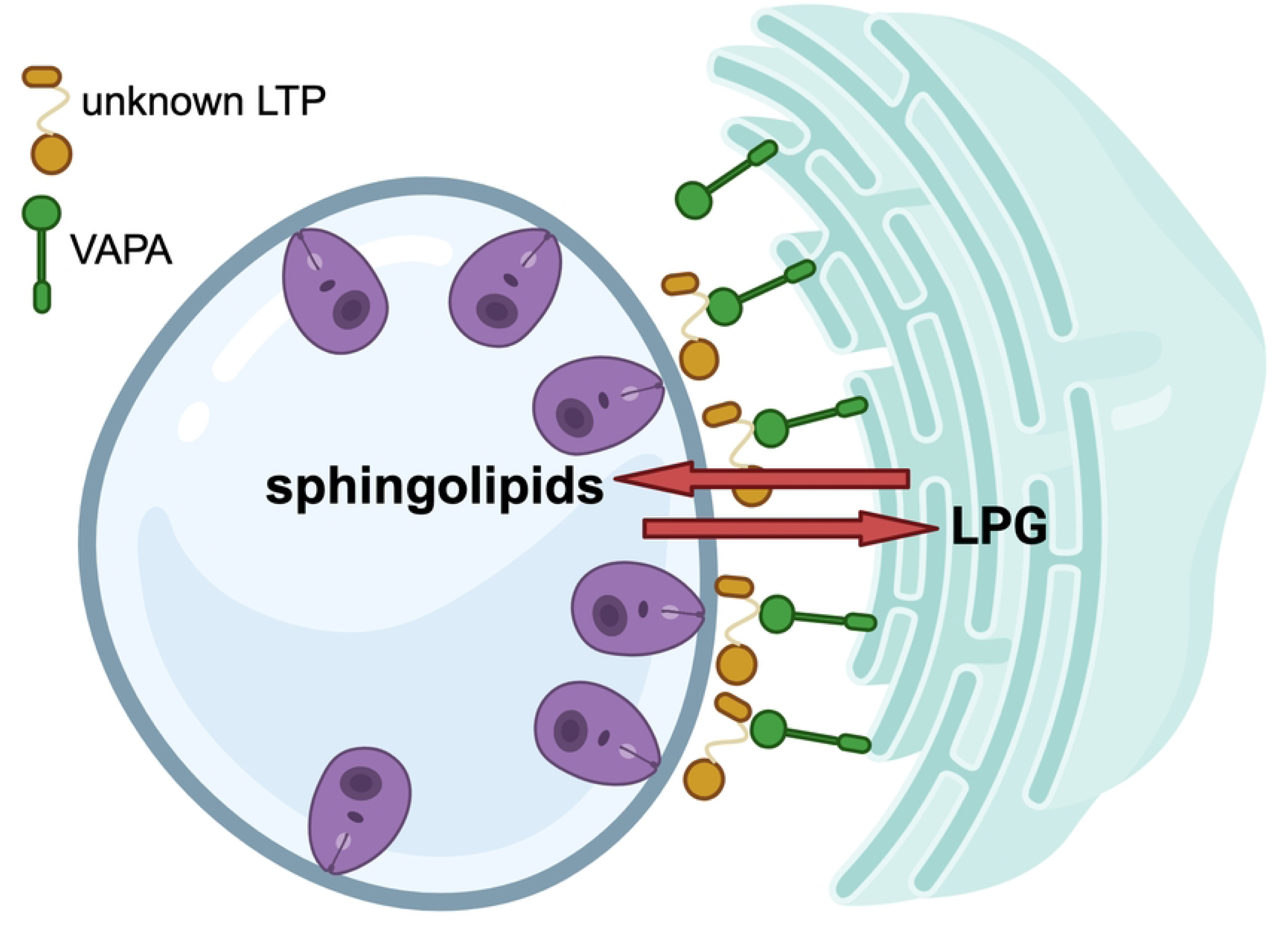
Proposed model for the role of VAPA in macrophages infected with *L. amazonensis*. VAPA is required for the acquisition of host sphingolipids by *L. amazonensis* amastigotes, as well as for the transfer of the *Leishmania* virulence glycolipid lipophosphoglycan (LPG) to the host cell endoplasmic reticulum. The identity of the lipid transfer protein(s) (LTP) involved in this bi-directional lipid transfer remains to be determined.

The vast majority of a cell’s organelles are connected to the ER through MCS, forming a network through which lipids, ions, and metabolites are travelling (30, 32, 51, 52). Our finding that VAPA accumulates around *L. amazonensis*-harboring PVs indicated that these PVs are connected through the ER to the host cell organelle network. Hence, one can envision that integration of *L. amazonensis*-harboring PVs in this ER-orchestrated organelle network facilitates the acquisition of host molecules by the parasite as well as the release of *Leishmania* molecules into the organellar network, and that these exchanges of molecules are essential for the ability of *L. amazonensis* to colonize macrophages. The observation that VAPA is required for the replication of *L. amazonensis* and for PV expansion supports the notion that contact between PVs and the ER plays a central role in the pathogenesis of *Leishmania* and is in agreement with a key role for the ER in this host-pathogen interaction (15, 41). It is also consistent with the notion that VAPA contributes to the replication of *Leishmania* by regulating the transfer of lipids from the ER to the PV.

In contrast to promastigotes, amastigotes are unable of *de novo* synthesis for most lipids and therefore rely on the salvage and remodeling of host lipids to satisfy their lipid requirements. (23, 25). In spite the fact that recent studies have investigated how *Leishmania* acquires host lipids, the underlying mechanisms remain poorly understood. Hence, a study on the role of the V-ATPase subunit ATP6V_0_d2 in the biogenesis of *L. amazonensis*-harboring PVs revealed its importance in PV enlargement (53). The authors of that study proposed that ATP6V_0_d2 participates in cholesterol influx, impacting the biogenesis of host cell membranes and PV formation (53), but cholesterol uptake by *L. amazonensis* was not directly evaluated. The discovery that the scavenger receptor CD36, a fatty acid transporter (54), accumulates on expanding PVs containing *L. amazonensis* and is required for parasite growth and PV expansion led the authors to suggest that CD36 may function as a channel to exchange lipids to and/or from the PV (55). In this regard, a potential source of fatty acids are lipid bodies (LBs), which accumulate in close proximity of PVs in *Leishmania*-infected macrophages (56, 57). Interestingly, a recent study shed light on how *L. donovani* may exploit LBs to acquire fatty acids (58). It was shown that biogenesis of LBs and their recruitment to *L. donovani*-containing PVs is mediated by Rab18 and TRAPPC9. Furthermore, the authors provided evidence that fluorescently-labelled fatty acids present in LBs were recruited in/on PVs containing *L. donovani* (58), suggesting that this parasite acquires fatty acids from PV-associated LBs. There was however no evidence in that study indicating that fluorescent fatty acids are incorporated by *Leishmania*. Sphingolipids received a particular attention as they are acquired and remodeled at high levels by *Leishmania* amastigotes (25) and are likely to play a key role in virulence (21, 23, 26, 59, 60). However, the mechanism by which *Leishmania* salvages sphingolipids remains elusive (23). Our results obtained using BODIPY FL C5-Ceramide add novel insight into this process, with VAPA being essential for the acquisition of fluorescent sphingolipids by *L. amazonensis* present in communal PVs. This approach has some limitations, since the use of BODIPY FL C5-Ceramide is not quantitative and it is not possible to determine the modifications that may have occured in infected macrophages. The identity of the fluorescent sphingolipid(s) detected in *L. amazonensis* therefore remains to be established. Given the importance of sphingolipids in cellular functions (61) and the essential role of VAPA in their acquisition by *L. amazonensis* amastigotes, it is reasonable to propose that the defect in parasite replication and PV expansion in the absence of VAPA is related to some extent to the inability to salvage host sphingolipids. A better understanding of how VAPA contributes to the acquisition of sphingolipids by *L. amazonensis* will be necessary to develop novel therapeutic approaches to interfere with the transfer of host sphingolipids to *Leishmania*.

The surface glycolipid LPG (48) plays a central role during the life cycle of *Leishmania*, as it contributes to the colonization of both the sand fly vector and the mammalian host (62–64). In macrophages, LPG enables *Leishmania* to interfere with signalling pathways (3, 65, 66), to inhibit phagolysosomal biogenesis (2, 4, 7), to activate the inflammasome (67), and to alter mitochondrial biology (68). To exert its action, LPG must notably exit the PV and redistribute within infected macrophages. We recently reported that upon phagocytosis, LPG is transported beyond the PV into the host cell ER and ERGIC via a mechanism dependent on the SNAREs Sec22b and syntaxin-5 (49). These findings are consistent with the notion that vesicular trafficking plays an important role in the spread of LPG inside infected macrophages. Interestingly, our results suggest that direct interaction between the PV and the ER through VAPA may also play a role in the transfer of LPG between these two organelles. Whether the Sec22b-dependent and the VAPA-dependent mechanisms involved in the transfer of LPG from the PV to the ER are part of the same pathway or represent two distinct pathways is not known. Interestingly, it has been reported that VAPA binds to and colocalizes with several SNAREs, including rsec22, the rat homologue of yeast sec22p (69). In yeast, Sec22 was shown to interact with lipid transfer proteins, and inhibition of Sec22 leads to defects in lipid metabolism at contact sites between the ER and plasma membrane (70). More recently, it was shown that Sec22b resides in phagosomal MCS where it acts as a thether at the ER-phagosome contacts and regulates phagosomal levels of phospholipids (71). Clearly, additional studies will be required to further investigate the mechanism(s) by which VAPA and Sec22b contribute to the transfer of LPG from the PV to the host cell ER and to identify the putative lipid transfer protein involved in this process. Whereas the impact of LPG on macrophage functions has been well described, the consequences of its presence specifically within the host cell ER remain to be established. ER stress induced by pathological conditions have been associated to the activation of the NLRP3 inflammasome (72). Previous work revealed that induction of ER stress in macrophages by *Leishmania* is associated with an increase survival of the parasite (73). Whether LPG plays a role in this process is unknown. In this regard, activation of the NLRP3 inflammasome by *Leishmania* is mediated by LPG and it requires that LPG be delivered out the PV (67). One may thus speculate that transfer of LPG from the PV to the host cell ER contributes to the ability of *Leishmania* to colonize macrophages, and that the impaired replication of *L. amazonensis* in VAPA knockdown macrophages may be in part related to the fact that LPG is not transferred to the ER. Future studies will be necessary to determine whether in addition to LPG, abundant glycolipids produced by *Leishmania* amastigotes such as glycoinositolphospholipids (74, 75) traffick to the host cell ER and whether VAPA plays a role in this process.

Lipid transport from the ER to the destination organelles requires the interaction of VAPA with various lipid transfer proteins, including CERT and ORP1L (42, 43). Whereas we observed by confocal immunofluorescence microscopy the presence of CERT and to a lesser extent ORP1L on PVs and their partial co-localization with VAPA, proximity ligation assays revealed that *L. amazonensis* efficiently disrupts the VAPA-CERT and VAPA-ORP1L *in situ* complexes present in BMM. The use of a *L. donovani* mutant deficient in LPG and a *L. major* mutant deficient in GP63 allowed us to discard a role for these two major pathogenicity factors in the disruption of VAPA-ORP1L complexes. However, we noted an increase in the number of those *in situ* complexes in the absence of GP63 at later time points post-infection, raising the possibility that GP63 plays a role in keeping VAPA from associating to ORP1L. Beyond the impact of GP63 on the VAPA-ORP1L *in situ* complexes observed at later time points post-infection, the mechanism by which interaction between VAPA and two prominent lipid transfer proteins is disrupted remains to be elucidated. Clearly, the rapid disruption of VAPA-CERT and VAPA-ORP1L complexes induced by *L. amazonensis* is consistent with a hijacking of VAPA by the parasite and raises the possibility that infection leads to the formation of novel VAPA-containing complexes which may contain proteins of either macrophage or parasite origin, or both. Such a scenario has been demonstrated in *Chlamydia*-infected cells where CERT is recruited to the *C. trachomatis* inclusion through interaction with the *Chlamydia* effector protein IncD (37). In addition to the recruitment of CERT and VAPA to the inclusion, two host cell sphingomyelin synthases are recruited to the inclusion, leading to the generation of a sphingomyelin factory at the inclusion (38). Whether VAPA forms complexes with a *Leishmania*-derived ceramide/sphingolipid transfer protein or associates with a sphingomyelin synthase as previously described for *Chlamydia* are issues that will deserve further investigation. Although our data suggest that CERT and ORP1L are not associated to VAPA in *L. amazonensis*-infected macrophages, we obtained evidence that both lipid transfer proteins play a role in the ability of *L. amazonensis* to replicate in macrophages. Hence, both CERT and ORP1L contribute to PV expansion, which may be related to the fact that sphingolipids and cholesterol are crucial components of cellular membranes (61, 76). It is also possible that absence of ORP1L results in the accumulation of damage to PV membrane, thereby preventing PV expansion (77). The observation that knockdown of ORP1L led to an increased replication of *L. amazonensis* might be related to the fact that absence of ORP1L may favor the interaction of VAPA with another lipid transfer protein. In this regard, it is important to consider that in addition to VAPA, the VAP family includes VAPB, MOSPD1, MOSPD2, and MOSPD3 (42). Over 250 human proteins were reported to interact with VAPA and VAPB and it was shown that these two VAPs share 50% of their interactomes (43), with the majority of these proteins being involved in lipid transfer between organelles (43). The fact that VAPA knockdown is sufficient to inhibit *L. amazonensis* replication and acquisition of sphingolipids suggests that there is no functional redundancy at that level within the VAP family. Whether other VAP family members are involved in other aspects of the interaction between *Leishmania a*nd macrophages will deserve further investigation.

In sum, we have provided evidence that VAPA, a component of MCS involved in the transfer of lipids between the ER and other organelles (42, 43), plays an important role in the ability of *L. amazonensis* to colonize macrophages. Integration of *L. amazonensis*-harboring communal PVs into the macrophage ER-orchestrated organelle network is consistent with the notion that the macrophage ER plays central role for the development of intracellular *Leishmania*.

## MATERIAL AND METHODS

### Ethic statement

Animal work was performed as stipulated by protocols 2112-01 and 2110-04, which were approved by the *Comité Institutionnel de Protection des Animaux* of the INRS-Centre Armand-Frappier Santé Biotechnologie. These protocols respect procedures on animal practice promulgated by the Canadian Council on Animal Care, described in the Guide to the Care and Use of Experimental Animals.

### Bone marrow-derived macrophages

Bone marrow was extracted from the femurs and tibias of 8 to 12 week-old female and male 129/BL6 mice. BMMs were differentiated for 7 days in complete Dulbecco’s modified Eagle’s medium with glutamine (DMEM; Thermo Fisher Scientific) containing 10% heat-inactivated fetal bovine serum (FBS; HyClone), 10 mM HEPES, pH 7.4, penicillin (100 IU/mL) and streptomycin (100 μg/mL) and supplemented with 15% (vol/vol) L929 cell-conditioned medium as a source of colony-stimulating factor-1 (CSF-1), in a 37°C incubator with 5% CO_2_. BMM were made quiescent by culturing them in DMEM without CSF-1 for 24 h prior to use.

### Parasite culture and infection

The *Leishmania* strains used in this study were *L. amazonensis* LV79 strain (MPRO/BR/72/M1841, obtained from the American Type Culture Collection), *L. donovani* LV9 (MHOM/ET/67/Hu3:LV9) Δ*lpg1* mutant and its complemented counterpart Δ*lpg1*+*LPG1* (78), and *L. major* NIHS (MHOM/SN/74/Seidman, obtained from Dr. W. Robert McMaster, University of British Columbia) Δ*gp63* mutant and its complemented counterpart *L. major* Δ*gp63*+*GP63* (79). Promastigotes were grown in *Leishmania* medium (M199 medium supplemented with 10% heat-inactivated fetal bovine serum [Multicell], 100 μM hypoxanthine, 10 mM HEPES, 5 μM hemin, 3 μM biopterin, 1 μM biotin, penicillin [100 U/ml], and streptomycin [100 μg/ml]) at 26°C. The *L. donovani* Δ*lpg1* +*LPG1* was cultured in *Leishmania* medium supplemented with zeocin (100 μg/mL) and the *L. major* Δ*gp63*+GP63 was cultured in *Leishmania* medium supplemented with G418 (100 μg/ml) to maintain the episomes. For infections, metacyclic promastigotes were enriched from late stationary-phase cultures using Ficoll gradients, as previously described (80). Complement opsonization of metacyclic promastigotes and zymosan was performed prior to macrophage internalization through incubation in Hank’s balanced salt solution (HBSS) containing 10% C5-deficient serum from female DBA/2 mice for 30 min at 37°C. Adherent BMM were then incubated at 37°C with metacyclic promastigotes or zymosan, and after 1h of incubation, noninternalized parasites were removed by washing three times with warm DMEM. Intracellular parasitemia was assessed at the indicated time point by counting the number of parasites per 100 infected BMM upon staining with the Hema 3 staining kit.

### Antibodies

The mouse anti-VAPA monoclonal antibody (MABN361) was from Millipore, the rabbit anti-CERT polyclonal antibody (PA5-115035) was from Invitrogen, the rabbit anti-ORP1L polyclonal antibody ORP1L was from Abcam (ab131165), the mouse anti-phosphoglycan (Galβ1,4Manα1-PO4) CA7AE monoclonal antibody (81) was from Cedarlane, the rabbit anti-β-actin polyclonal antibody was from Cell Signalling, and the rat LAMP-1 monoclonal antibody 1D4B developed by J.T.August and purchased through the Developmental Studies Hybridoma Bank at the University of lowa and the National Institute of Child Health and Human Development. The secondary antibody anti-mouse conjugated to Alexa Fluor 488, and the secondary antibody anti-rabbit conjugated to Alexa Fluor 568 and the secondary antibody anti-rat conjugated to Alexa Fluor 647 used for immunofluorescence were from Invitrogen-Molecular probes.

### Small interfering RNA knockdown

The protocol used for siRNA transfection was adapted from Dharmacon’s cell transfection (www.dharmacon.com). BMM were seeded onto 24-well plates and reverse-transfected with the Lipofectamine RNAiMAX Reagent (Invitrogen) as per the manufacturer’s recommendation. The final concentration of siRNA was 80 nM in a final volume of 200 μl of complete DMEM. BMM were either mock-transfected, transfected with the Non-Targeting siRNA (Dharmacon) with the following target sequence: UAGCGACUAAACACAUCAA, or transfected with the following ON-TARGET SMARTpool siRNA: Vapa (Dharmacon), which contains four siRNA with the following target sequences: sequence 1, CCAUCGGAUAGAAAAGUGU; sequence 2, CAGCCAUUUUCAUUGGAUU; sequence 3, CUACAAAUCUUAAAUUGCA; sequence 4, GUGUUUCACUCAAUGAUAC; Osbp (Dharmacon) which contains four siRNA with following target sequences: sequence 1, GCGAUGAAGAUGACGAGAA; sequence 2, AUGAAAGAGACCAGCGAAU; sequence 3, GAGAAUGGGUAUCGGUCCA; sequence 4, GAAGAGGGCUGAUCGGAGA; Col4a3bp (Dharmacon) which contains four siRNA with following target sequences: sequence 1, UCAGAGGGAUAAAGUCGUA; sequence 2, GCGAGAAUACCCUAAGUUU; sequence 3, UCAGAGAGACGUACUGUAU; sequence 4, GAUCACGUAUGUAGCUAAU. After 48 h cells were washed with complete medium prior to being used.

### BODIPY FL-C5-Ceramide labeling and live cell microscopy

Controls and siRNA to VAPA-treated BMM seeded onto glass coverslips and infected for 48 h were washed three times with cold Hank’s Balanced Salt Solution (HBSS, Invitrogen) and incubated in DMEM (without FBS) containing 0.625 μM of BODIPYFL-C5-Ceramide (Invitrogen) for 30 min at 4°C. The cells were then washed three times with cold HBSS and incubated in DMEM without FBS at 37°C in the presence of 5% CO_2._ Images of live cells were acquired in the channel 489-569 nm using a LSM780 confocal microscope (Carl Zeiss Microimaging) using X 63 objective every 5 min after the beginning of the chase.

### Confocal immunofluorescence microscopy

BMM were seeded in 24-well plates containing microscope coverslips and were infected with *Leishmania* metacyclic promastigotes as described above or were fed zymosan for the indicated time points. Cells were washed with phosphate-buffered saline (PBS), fixed with 3.7% paraformaldehyde (PFA) for 30 min, and then permeabilized in 0.1% Triton X-100 for 5 min. Samples were next blocked in 10% bovine serum albumin for 1 h. Cells were incubated for 1 h the various antibodies at the following dilutions: 1:400 for the mouse anti-VAPA monoclonal antibody, 1:400 for the rabbit anti-CERT polyclonal antibody, 1:200 for the rabbit anti-ORP1L polyclonal antibody, and 1:200 for the CA7AE mouse monoclonal antibody. BMM were next incubated for 1 h with an appropriate combination of secondary antibodies (1:500 for the anti-mouse Alexa Fluor 488; 1:500 for the anti-rabbit Alexa Fluor 568, and 1:500 for the anti-IgM Alexa Fluor 568. Macrophage and promastigote nuclei were stained with DAPI (Molecular Probes). Coverslips were washed three times with PBS after every step, and individually mounted with Fluoromount-G (Invitrogen) onto slides for imaging under a confocal microscope. Analyses of distribution were performed on a LSM780 confocal microscope (Carl Zeiss Microimaging) using Plan Apochromat x 63 oil-immersion differential interference contrast (DIC) (NA 1.64) objective, and images were acquired in sequential scanning mode. Images were processed with ZEN 2012 software. At least 25 cells per condition were analyzed using the Icy image analysis software or ZEN 2012 software, and statistical differences were evaluated using one-way analysis of variance (ANOVA) followed by Tukey’s posttests (three groups). Data were considered statistically significant when *P* was <0.05, and graphs were plotted with GraphPad Prism 5.

### Proximity ligation assay

Visualization of *in situ* protein-protein interactions was carried out according to the manufacturer’s protocol using Duolink *In Situ* Detection Reagents (Sigma Aldrich, DUO92004, DUO92002, DUO92008). BMM were seeded onto glass coverslips in 24-well plates and infected with *Leishmania* metacyclic promastigotes for the indicated time points. Cells were washed with phosphate-buffered saline (PBS) and fixed with 3.7% PFA for 30 min. Cells were next permeabilized with 0.2% Triton X-100. Blocking was (Sigma Aldrich, DUO92004) followed by 2 h incubations with primary antibodies directed either against VAPA (Millipore, 1:200), CERT (Invitrogen, 1:400) or ORP1L (Abcam, 1:200). After incubation the primary antibodies were washed away and PLA secondary probes were added; Anti-Rabbit MINUS (Sigma Aldrich, DUO92004, dilution 1:5), and Anti-Mouse PLUS (Sigma Aldrich, DUO92002, dilution 1:5) following by 1 h incubation in a humidified chamber at 37°C. The coverslips were twice washed in Duolink *In situ* wash buffer A for 5 min under gentle agitation. Ligation was then done by adding Ligation buffer (Sigma Aldrich, dilution 1:5) and Ligase (Sigma Aldrich, dilution 1:40) solutions to each sample and for 30 min incubation in a humidified chambre at 37°C. The coverslips were then washed twice in 1X Duolink In Situ wash buffer A for 2 min under gentle agitation, and then placed in amplification buffer (Sigma Aldrich, DUO92008,dilution 1:80) solution and incubated 100 min at 37°C, followed by washing twice in 1X wash buffer B (Sigma Aldrich) and then once for 1 min in 100 X diluted wash buffer B, and drying at room temperature in the dark. Coverslips were incubated for additional 10 min with DAPI and then individually mounted with Fluoromount-G (Invitrogen, ref.00-4958-02) onto slides. Analyses were performed on a LSM780 confocal microscope (Carl Zeiss Microimaging) using Plan Apochromat X 63 oil-immersion differential interference contrast (DIC). To determine the background PLA signal, the same procedures were followed as described above except that the cell-containing coverslips were incubated with only one of the individual primary antibodies. Images of PLA are visualized as red dots and were analyzed with Zen software by manual counting. The imaging results are presented as the mean of the three trial values ± standard deviation. Statistical significance (*p*-values) was determined using a one-way ANOVA test in the software GraphPad Prism 5.

### Western blot analyses

Prior to lysis, adherent BMMs were placed on ice and washed 3 times with PBS containing 1 mM sodium orthovanadate and 5 mM 1,10-phenanthroline (47). Cells were scraped in the presence of lysis buffer containing 1% NP-40, 50 mM Tris-HCl (pH 7.5), 150 mM NaCl, 1 mM EDTA (pH 8), 10 mM 1,10-phenanthroline, and phosphatase and protease inhibitors (Roche) (47). After 24 h incubation at −20°C, lysates were centrifuged for 30 min, and the soluble phase was collected.

After protein quantification, 25 μg of protein was boiled (100°C) for 5 min in SDS sample buffer and migrated in SDS-PAGE gels. Proteins were transferred onto Hybond-ECL membranes (Amersham Biosciences), blocked for 1 h in Tris-buffered saline (TBS) 1× 0.1% Tween containing 5% BSA, and incubated with primary antibodies (diluted in TBS 1× 0.1% Tween containing 5% BSA) overnight at 4°C and then with appropriate horseradish peroxidase (HRP)- conjugated secondary antibodies for 1 h at room temperature. Membranes were incubated in ECL (GE Healthcare), and immunodetection was achieved via chemiluminescence using the Azure Biosystem.

### Statistics and reproducibility

GraphPad Prism 5 software was used to generate the graphs and statistical analyses. All experiments were conducted three independent times. Methods for statistical tests, exact value of *n*, and definition of error bars are indicated in the figure legends; *, *P* < 0.05; **, *P* < 0.01; ***, *P* < 0.001. All immunoblots and images shown are representative of these independent experiments.

## ACKNOWLEDGMENTS

We thank Dr Laurent Chatel-Chaix and Dr Olus Uyar for advice with proximity ligation assays, Dr Acevedo Ospina for thoughtful comments on this manuscript, and Mr. Jessy Tremblay for expert assistance in immunofluorescence confocal microscopy experiments.

## AUTHOR CONTRIBUTIONS

Conceptualization, I.G. and A.D.; Methodology, I.G. and A.D.; Formal Analysis, I.G. and A.D.; Investigation, I.G.; Resources, A.D.; Writing - Original Draft, I.G. and A.D.; Writing - Review and Editing, A.D.; Visualization, I.G. and A.D.; Supervision, A.D.; Funding Acquisition, A.D.

